# SARS-CoV-2 and Malayan pangolin coronavirus infect human endoderm, ectoderm and induced lung progenitor cells

**DOI:** 10.1101/2020.09.25.313270

**Authors:** Bixia Hong, Xinyuan Lai, Yangzhen Chen, Tianming Luo, Xiaoping An, Lihua Song, Hui Zhuang, Huahao Fan, Tong Li, Kuanhui Xiang, Yigang Tong

**Affiliations:** Beijing Advanced Innovation Center for Soft Matter Science and Engineering, College of Life Science and Technology, Beijing University of Chemical Technology, Beijing 100029, China; Department of Microbiology and Infectious Disease Center, School of Basic Medical Sciences, Peking University Health Science Center, Beijing 100191, China

**Keywords:** Coronavirus disease 19, Severe acute respiratory syndrome coronavirus 2, pangolin coronavirus, human embryonic stem cells, stem cell differentiation

## Abstract

Since the infection of severe acute respiratory syndrome coronavirus 2 (SARS-CoV-2) in several somatic cells, little is known about the infection of SASRS-CoV-2 and its related pangolin coronavirus (GX_P2V). Here we present for the first time that SARS-CoV-2 pseudovirus and GX_P2V could infect lung progenitor and even anterior foregut endoderm cells causing these cells death, which differentiated from human embryonic stem cells (hESCs). The infection and replication of SARS-CoV-2 and GX_P2V were inhibited when treated with whey protein of breastmilk and Remdesivir, confirming that these two viruses could infect lung progenitor and even anterior foregut endoderm. Moreover, we found that SARS-CoV-2 pseudovirus could infect endoderm and ectoderm. We found that whey protein blocked SARS-CoV-2 infecting these cells. In line with the SARS-CoV-2 results, GX_P2V could also infected endoderm and ectoderm, and also was inhibited by Remdesivir treatment. Although expressing coronavirus related receptor such as ACE2 and TMPRSS2, mesoderm cells are not permissive for SARS-CoV-2 and GX_P2V infection, which needed further to study the mechanisms. Interestingly, we also found that hESCs, which also express ACE2 and TMPRSS2 markers, are permissive for GX_P2V but not SARS-CoV-2 pseudovirus infection and replication, indicating the widespread cell types for GX_P2V infection. Heparin treatment blocked efficiently viral infection. These results provided insight that these stem cells maybe provided a stable repository of coronavirus function or genome. The potential consequence of SARS-CoV-2 and animal coronavirus such as GX_P2V infection in hESCs, germ layer and induced progenitors should be closely monitored.

## Introduction

Coronavirus disease 2019 (COVID-19) caused by the novel coronavirus, SARS-CoV-2, was declared a pandemic by World Health Organization (WHO) and resulted in over 30 million confirmed cases and more than 800 thousand deaths over the world since its outbreak. Although respiratory failure is the most common mortality outcome in COVID-19 patients, other fatal manifestations are also existed in multiple organ systems, including the heart, brain and gastrointestinal tract^1^. It’s reported that nearly 25% of the 400 COVID-19 patients had suffered in-hospital cardiac injury ^2^. Nearly 25% of the COVID-19 patients had gastrointestinal symptom, including anorexia, diarrhea, vomiting and abdominal pain, indicating that SARS-CoV-2 may infects gastrointestinal cells ^3^. Interestingly, a recent Germany study demonstrated that positive SARS-CoV-2 RNA were detected in human brain biopsies in 36.4% of fatal COVID-19 cases ^4^. Recently, a group found that SARS-CoV-2 pseudovirus could infect pancreatic endocrine cells, liver organoids, cardiomyocytes and dopaminergic neurons derived from human pluripotent stem cells (hPSCs) ^5^. All these findings highlight the potential of viral infection in several organs and even stem cells. However, there has been no direct experimental evidence of SARS-CoV-2 infection in different differentiation stages of human embryonic stem cells. A recent clinical study showed that SARS-CoV-2 related receptors, such as ACE2 and TMPRSS2, also express in the trophoblast of the blastocyst and tiotrophoblast and hypoblast of the implantation stages, indicating that the embryo may support SARS-CoV-2 infection resulting potential risks for vertical transmission and fetal health ^6^. Importantly, a recent study showed that SARS-CoV-2 could also infect human neural progenitor cells, indicating that SARS-CoV-2 could infect and replicate in widespread cell types even stem cells. Together, these findings suggest that the different stages of human embryonic stem cells differentiation might also be the susceptible for SARS-CoV-2 infection.

## Materials and Methods

### Maintenance of hESCs

hESCs were maintained in mTeSRTM1 medium (STEMCELL Technologies, Vancouver, Canada) as we previously described with daily medium replenishment. Cells were passaged with ReleSR (STEMCELL Technologies) when cells reach approximately 80% confluence. Cells were cultured in a 5% CO_2_ air environment.

### Other cell lines, coronavirus and key reagents

Vero E6 cells were cultured in high-glucose-containing Dulbecco’s Modified Eagle Medium supplemented with 10% fetal bovine serum. GX_P2V was described recently^7^. SARS-CoV-2 pseudovirus was kindly shared by Prof. Youchun Wang (National Institutes for Food and Drug Control, China)^8^. Goat and cow whey proteins were purchased from Sigma (USA). Heparin was purchased from Sigma (USA). Remdesivir was purchased from Sigma (USA).

### Stem cell differentiation

#### Endoderm

The hESCs were differentiated into definitive endoderm using the commercial STEMdiff Definitive Endoderm kit (STEMCELL Technologies) according to the manufacturer’s instructions. Briefly, hESCs were dissociated into single cells with Accutase and plated onto Matrigel coated plates. The next day, differentiation was induced by media A of the STEMdiff Definitive Endoderm kit for one day and then media B for more three days.

#### Mesoderm

mesoderm induced differentiation was performed by the STEMdiff Mesoderm induction Medium (STEMCELL Techonoloties) according to the manufacturer’s instructions. Briefly, dissociated hESCs cells were plated onto the Matrigel-coated (BD) plates with ROCK inhibitor of Y-27632 (STEMCELL Technologies) at the concentration of 10 μM. The next day, the medium was exchanged with STEMdiff Mesoderm induction Medium for four days, with daily medium replenishment.

#### Ectoderm

the neuroectoderm cells were induced by using the dual SMAD inhibition method as previously described in the literature with some modifications. Briefly, Single dissociated hESCs cells were plated on the Matrigel-coated plates with ROCK inhibitor (Y-27632). The next day, ectoderm differentiation was performed with basal Medium (DMEM/F12, 10% knockout serum replacement (KOSR) and 1% Glutamax) supplemented with 10 μM SB431542 (STEMCELL Technologies) and 200 nM Noggin (Peprotech) for four days.

### Induction of anterior foregut endoderm

For anterior foregut endoderm induction (hAFECs), the differentiation was performed as described previously with some modification^9^. Briefly, the endodermal cells were dissociated into single cells using Accutase and plated on the Matrigel-coated plates in serum-free differentiation media (SFM) supplemented with 1.5 μM Dorsomorphin dihydrochloride (R&D System) and 10 μM SB431542 for 24h. The next day, the media were exchanged with SFM supplemented with 10 μM SB431542 and 1 μM IWP2 (R&D System) for two days. SFM: DMEM/F12 supplemented with N2, B27, 50 μg/ml ascorbic acid, 2 mM Glutamax, 0.4 μM and 0.05% bovine serum albumin (BSA).

### Induction of lung progenitors

For human lung progenitor cells (hLPCs) induction, the induced hAFECs was cultured with the SFM medium supplemented with 3 μM CHIR99021 (STEMCELL Technologies), 10 ng/ml BMP4 (Peprotech), 20 ng/ml FGF10 (Peprotech), 10 ng/ml FGF7 (Peprotech) and all-trans retinoic acid (ATRA) (Sigma) for 8 days, with every other day medium replenishment.

### Viral infection assay

SARS-CoV-2 pseudovirus infection was performed as previously described^10^. The cells were infected with viral inocula of 650 TCID_50_/well^8^. One day post infection (1dpi), the cells were lysed and the luminescence was measured according to the manufacture’s protocol. Cells were infected with GX_P2V at multiplicity of infection (MOI) of 0.01 as described^11^. Remdsivir, heparin and human breastmilk were utilized to inhibit viral infection and replication during the infection process.

### mRNA quantification

GX_P2V RNA quantification was performed by RT-qPCR as previously described^11^. Briefly, total RNA was extracted by the AxyPrep™ multisource total RNA Miniprep kit (Axygene). Hifair II 1st Strand cDNA synthesis kit with gDNA digester (Yeasen) was used to synthesize the first strand complementary DNA (cDNA). Hieff qPCR SYBR Green Master Mix (Yeasen) was used to quantify GX_P2V RNA. The RT-qPCR amplification of the Taqman method was performed as follows: 50°C for 2 min, 95°C for 10 min followed by 40 cycles consisting of 95°C for 10 s, 60°C for 1 min. The markers related such as Sox2, FoxA2, NKx2.1, ACE2 and TMPRSS2 were quantified by RT-qPCR normalized by RPS11. The primers were shown in supplementary table 1.

### Western blotting

Western blotting was performed as described previously^12^. Antibodies against nucleocapsid protein of anti-SARS-CoV-2 N protein (Genscript) and GAPDH of anti-GAPDH (Proteintech) were used at 1:3000 dilutions. The second antibody of HRP-conjugated affinipure Goat anti-mouse IgG (H+L) was diluted at 1:20000. SuperSignal^®^ West Femto Maximum Sensitivity Chemiluminescent Substrate (Thermo Scientific, USA) was used for imaging.

### Immunofluorescent assay

Cells were fixed by 4% para-formadehyde in phosphate-buffered saline (PBS) at room temperature for 10 min and permeabilized by 0.1% Triton-100 at room temperature for 10 min. Then, the cells were blocked with blocking buffer (10% goat serum and 1% BSA in PBS) for 1hr. Cells were incubated with primary antibodies (diluted in blocking buffer) at 4°C overnight. The primary antibodies are anti-OCT4 rabbit polyclonal antibody (Proteintech), anti-SSEA-4 antibody (STEMCELL Technologies), anti-Sox17 antibody (R&D System), anti-Brachyury antibody (Cell Signaling Technology) and ani-Nestin antibody (R&D System). The Alexa-488 or 594 conjugated secondary antibodies (1:2000 diluted in blocking buffer) were added to the cells in the dark at room temperature for 1 hr. Nuclei were stained with DAPI for 10 min at room temperature.

### Plaque assay for determining virus titer

The plaque assay for determining virus titer was performed as previously described^11^. Briefly, confluent monolayer Vero E6 cells were infected with GX_P2V with ten-fold dilution from 10^−1^ to 10^−6^. After removing the virus, the cells were washed by PBS and added with 1% agarose overlay to prevent cross-contamination. At 5 dpi, cells were fixed with 4% paraformaldehyde for 1h, followed by staining with Crystal violet for 10 min and washed with water. The plaques were counted and virus titers were calculated.

### Statistical analysis

Statistical analyses were analyzed using GraphPad Prism 8 software (GraphPad Software Inc., San Diego, CA, USA). Values are shown as mean of triplicates. Comparisons between the two groups were analyzed using the Student’s t tests. Values of *p*<0.05 was considered statistically significant.

## Results

To investigate whether SARS-CoV-2 infects human lung progenitor cells (hLPCs), we first evaluated the expression of ACE2 and TMPRSS2 in hLPCs. Our data suggested that both ACE2 and TMPRSS2, as well as lung progenitor markers of Sox2, FoxA2 and NKx2.1, were detected in hLPCs (**Fig. S1A and C**). Then, we infected the hLPCs with SARS-CoV-2 pseudovirus at viral inocula of 650 TCID_50_/well as previously described^13^. As a result, SARS-CoV-2 pseudovirus could infect hLPCs and the infection could be inhibited by breastmilk (2 mg/ml), which was reported to have anti-SARS-CoV-2 activity ^13^(**Fig. 1A**). In line with the SARS-CoV-2 pseudovirus results, we also infected hLPCs with SARS-CoV-2 related pangolin coronavirus (GX_P2V) ^7^at 0.1 multiplicity of infection (MOI) and found that GX_P2V RNA could be detected in hLPCs^11^. After Remdesivir (1.5 μM) and human breastmilk (2 mg/ml) treatment, GX_P2V RNA level decreased significantly compared to the control (**Fig. 1B**). In addition, we also analyzed the cell viability of GX_P2V infected hLPCs. Accordingly, GX_P2V infection reduced the viability of hLPCs. Next, we challenged the earlier stage cells of human anterior foregut endoderm (hAFECs) with SARS-CoV-2 pseudovirus (650 TCID_50_/well) and GX_P2V (MOI=0.1) for viral infection assessment. We found that hAFECs markers and also ACE2 and TMPRSS2 expressed in the differentiated cells (**Fig. S1B**). Luciferase activity could be detected and decreased after breastmilk (from both human and goat) treatment in hAFECs (**Fig. 1C**). Interestingly, GX_P2V RNA levels decreased during the Remdesivir (1.5 μM) and human breastmilk (2 mg/ml) treatment in the hAFECs (**Fig. 1D**). In addition, we also found that heparin could also significantly inhibited SARS-CoV-2 pseudovirus infection (**Fig. S2A-C**), indicating heparins potential for treatment of SARS-CoV-2 infection, and the heparan sulphate could be the potential co-receptor for SARS-CoV-2 infection^14^. These findings suggest that hLPCs and also hAFECs are permissive for SARS-CoV-2 pseudovirus and GX_P2V infection.

**Fig. 1.**
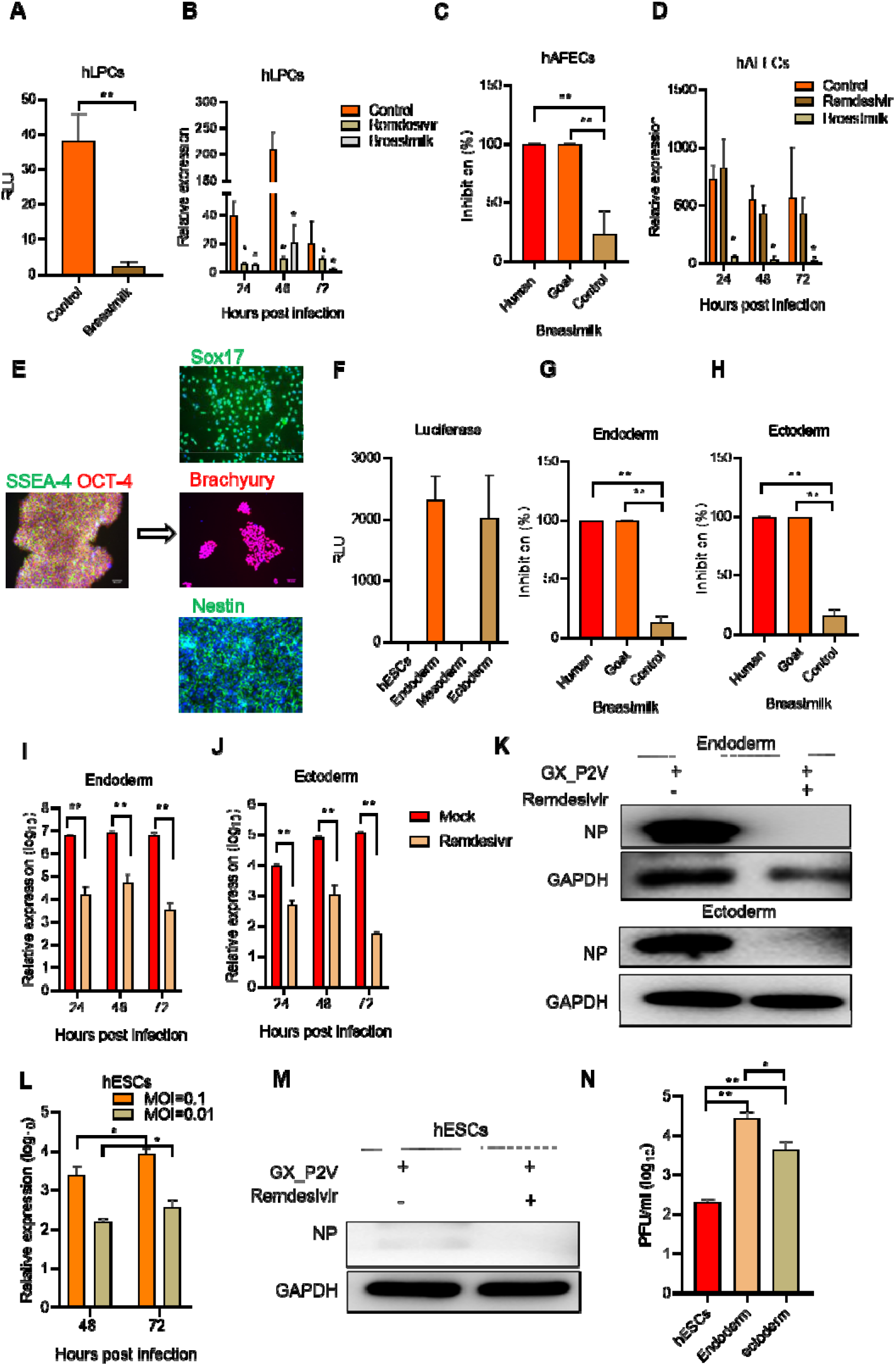
SARS-CoV-2 pseudovirus and GX_P2V infect hESCs, endoderm, ectoderm, hAFECs and hLPCs. **(A)** hLPCs and anterior foregut endoderm **(C)** were infected with SARS-CoV-2 pseudovirus at viral inocula of 650 TCID50/well with or without breastmilk (2 mg/ml). Cell were harvest at 24 hpi and luciferase activity was determined. **(B)** hLPCs and hAFECs **(D)** were challenged with 0.1 MOI GX_P2V combined with Remdsivir (1.5 μM) and human breastmilk (2mg/ml), respectively. Cells were harvested at 24, 48 and 72 hpi and viral loads were quantified by RT-qPCR**. (E)** Representative images of hESCs markers of SSEA4 andOCT-4, endoderm marker of Sox17, mesoderm marker of Brachyury and ectoderm marker of Nestin. Scale bars, 100 μm. **(F)** SARS-CoV-2 (650 TCID50/well) infection was performed in hESCs, endoderm, mesoderm and ectoderm and luciferase activity was detected at 24 hpi. Human breastmilk of human and goat were used to treat SARS-CoV-2 pseudovirurs infection in endoderm **(G)** and ectoderm **(H).** Endoderm **(I)** and ectoderm **(J)** were infected with GX_P2V (MOI 0.1) and treated with Remdesivir (1.5 μM) and viral loads were determined by RT-qPCR at 24, 48 and 72 hpi. **(K)** Representative image of NP of GX_P2V expression in endoderm and endoderm cells. hESCs were challenged with GX_P2V (MOI 0.1 and 0.01) and viral loads were detected by RT-qPCR at 48 and 72 hpi (**L**) and by western blot at 72 hpi **(M). (N)** Infectious virus titer of the supernatant samples from hESCs, endoderm and ectoderm infected with GX_P2V for 72h were determined by plaque assay. Data were obtained from three independent experiments. hESCs, human embryonic stem cells; hAFECs, human anterior foregut endoderm; hLPCs, human lung progenitor cells; MOI, multiplicity of infection; hpi, hours post-infection; NP, nucleoprotein. Values are shown as mean of triplicates ± SD, *p<0.05, **p<0.01 by unpaired two-tailed t test.

To explore the possible permissive of endoderm, mesoderm and ectoderm to SARS-CoV-2 and GX_P2V infection, we generated hESCs into endoderm, mesoderm and ectoderm according to the manufacturer’s instructions, respectively. As shown in **Fig. 1E**, the immunofluorescent assay showed that Sox17, Brachyury, nestin was detected in endoderm, mesoderm and ectoderm, respectively. In addition, ACE2 and TMPRSS2 were also expressed in these cells (**Fig. S1C**). Next, we challenged these cells with SARS-CoV-2 pseudovirus and GX_P2V virus. As shown in **Fig. 1F**, SARS-CoV-2 pseudovirus could be detected in endoderm and ectoderm, but not mesoderm. The SARS-CoV-2 infection could be inhibited by whey protein of human and goat both in endoderm (**Fig. 1G**) and ectoderm (**Fig. 1H**). However, the cells activity from hESCs, endoderm and ectoderm was influenced by human breastmilk (data not shown). Interestingly, GX_P2V RNA was positive in endoderm and ectoderm cells but not mesoderm, indicating the potential inhibitory activity or lack key factors of mesoderm to SARS-CoV-2 and GX_P2V infection. After Remdesivir (1.5 μM) treatment, GX_P2V RNA level was significantly lower compared to those without any drug treatment both in endoderm (**Fig. 1I**) and ectoderm (**Fig. 1J**). We found that GX_P2V nucleoprotein (NP) could be detected in endoderm and ectoderm (**Fig. 1K**), but not in mesoderm (data not shown) by western blot assay. In addition, we also challenged the Vero E6 cells with the supernatant collected from infected endoderm and ectoderm to perform plaque assay. As results, both endoderm and ectoderm could reproduce infectious viruses (**Fig. 1N**). These results indicated that the germ layer cells such as endoderm and ectoderm are also permissive for both SARS-CoV-2 and GX_P2V infection.

To further investigate whether SARS-CoV-2 pseudovirus and GX_P2V infect hESCs, the cultured hESCs were challenged by these two viruses. Firstly, we detected the stem cell markers of OCT-4 and SSEA-4 by immunofluorescent assay. As shown in **Fig. 1E**, OCT-4 and SSEA-4 staining were identified in the hESCs cells. We infected the hESCs with SARS-CoV-2 pseudovirus and GX_P2V, respectively. Unfortunately, luciferase activity was undetected in the SARS-CoV-2 pseudovirus infected hESCs cells (**Fig. 1F**). However, GX_P2V RNA was relative positive at 48 and 72 hours post infection (hpi) (**Fig. 1L**) and NP was positive in hESCs cells (**Fig. 1M**). Remdesivir also showed inhibitory activity to GX_P2V in these cells. Plaque assay results showed that hESCs could also produce infectious GX_P2V. These results indicated that hESCs are not permissive for SARS-CoV-2 pseudovirus but permissive for the pangolin coronavirus of GX_P2V infection.

## Discussion

We demonstrated that induced endoderm, ectoderm, hAFECs and hLPCs are permissive for both SARS-CoV-2 and GX_P2V infection. The hESCs are permissive for GX_P2V but not SARS-CoV-2 pseudovirus infection. Extensive viral protein expression and infectious viral particles were also detected in these infected cells. Breastmilk and Remdesivir could inhibit viral infection and replication in these cells. In addition, heparin could also inhibit both SARS-CoV-2 and GX_P2V infection, indicating its potential usage as a direct-acting inhibitor of SARS-CoV-2 and GX_P2V infection. These results may explain why SARS-CoV-2 could infect many organs. Moreover, these results provided insight that these stem cells maybe provided a stable repository of coronavirus function or genome. Coronavirus persistence in human stem cells confers the possibility that coronavirus may eventually become a passenger in the human germline and even vertical transmission to new born. Surprisingly, the pangolin coronavirus of GX_P2V could not only infect human mature somatic cells^1^, but also infect the hESCs, endoderm, ectoderm, hAFECs and hLPCs, indicating the possibility of the cross-species transmission and potential danger of coronavirus from animals. However, both viruses could not infect mesoderm, even though it expresses viral recptor of ACE2 and TMPRSS2, indicating the potential inhibitory activity of mesoderm to these coronaviruses, which may explain why mesenchymal stem cells could be used as a therapy tool for COVID-19 treatment^15^. The potential consequence of SARS-CoV-2 and animal coronavirus such as GX_P2V infection in hESCs, germ layer and induced progenitors should be closely monitored.

## Contributions

K.X., Y.G., H.F. and T.Li. designed the research; H.F., B.H., X.L., YC., T.L., X.A. T. Luo, L.S. and Y.H. performed the experiments; K.X., and H.F. analyzed the data; K.X., H.F., T.Li, H.Z., and Y.T. wrote and revised the manuscript.

## Acknowledgements

We gratefully thank Prof. Youchun Wang and Weijin Huang (National Institutes for Food and Drug Control, China) for sharing the pseudovirus of SARS-CoV-2.

## Supplementary tables

**Supplementary Table 1:**
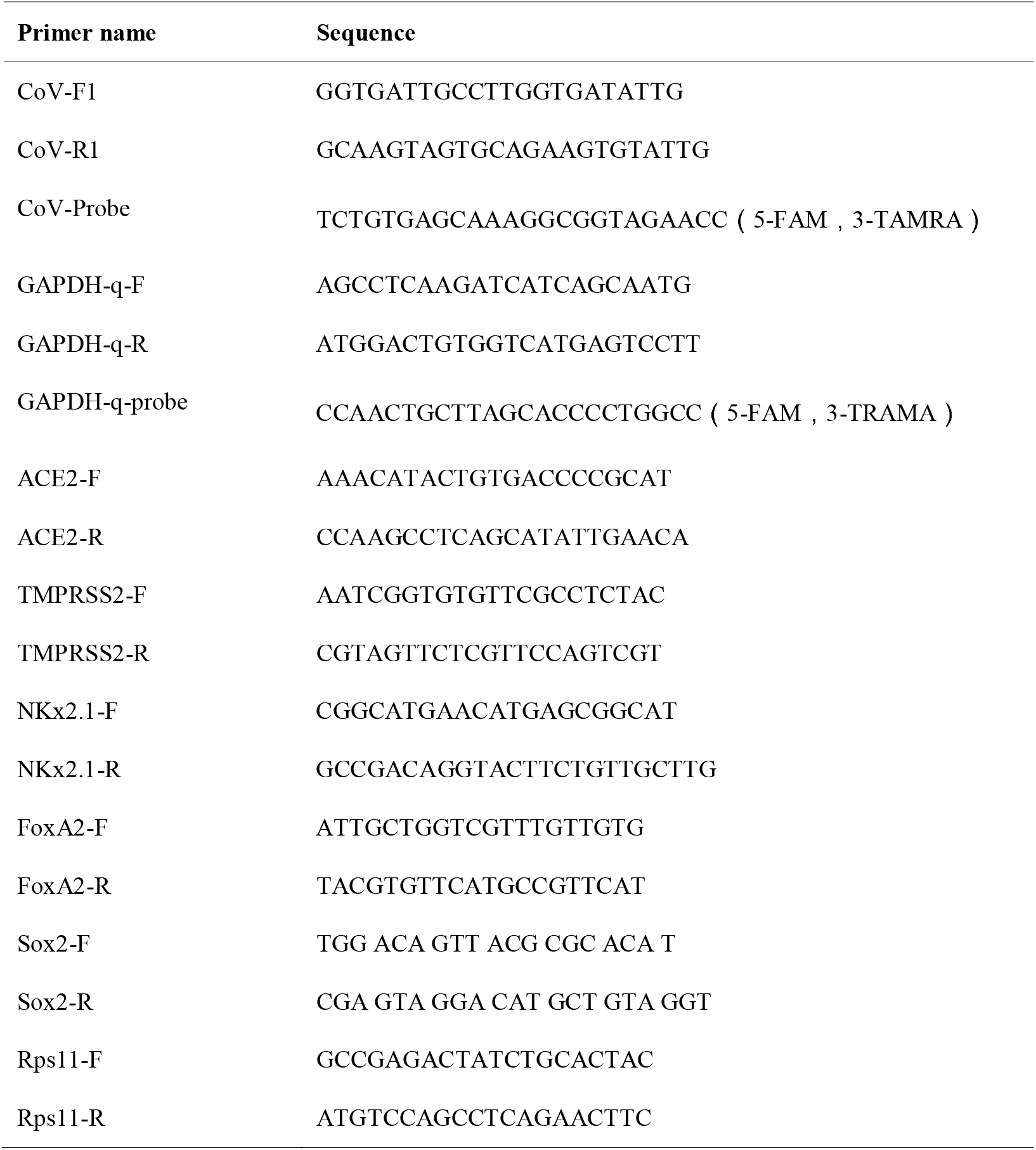
Primers used in the study.

## Supplementary figures and figure legends

**Fig. S1.**
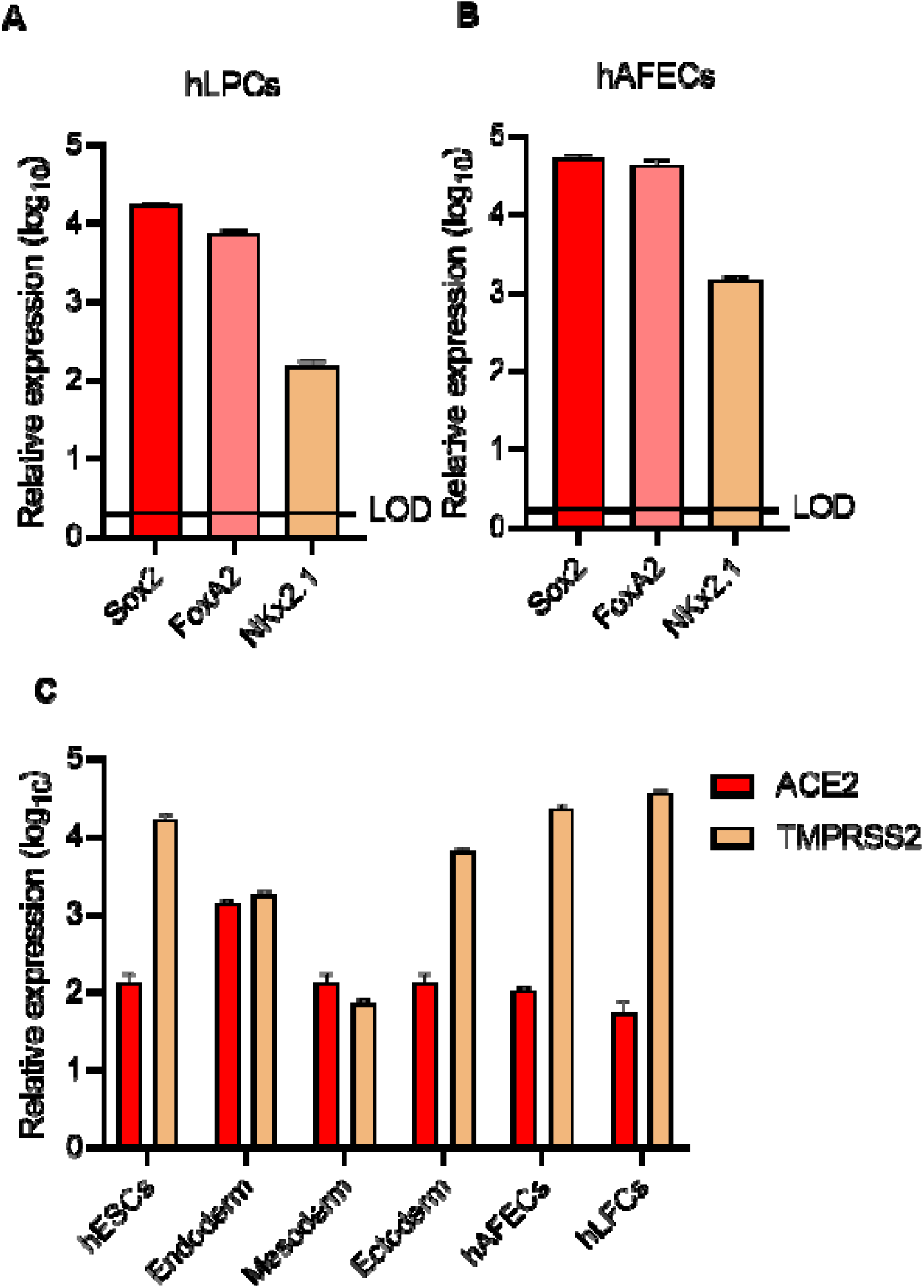
Lung progenitor related markers and coronavirus receptor expression in hESCs, endoderm, mesoderm, ectoderm, hAFECs and hLPCs. RT-qPCR analysis of endoderm and lung progenitor related markers of Sox2, FoxA2 and NKx2.1 in hLPCs (**A**) and hAFECs (**B**). (**C**) RT-qPCR analysis of coronavirus related receptor of ACE2 and TMPRSS2 expression in hESCs, endoderm, mesoderm, ectoderm, hAFECs and hLFCs. hESCs, human embryonic stem cells; hAFECs, human anterior foregut endoderm; hLPCs, human lung progenitor cells.

**Fig. S2.**
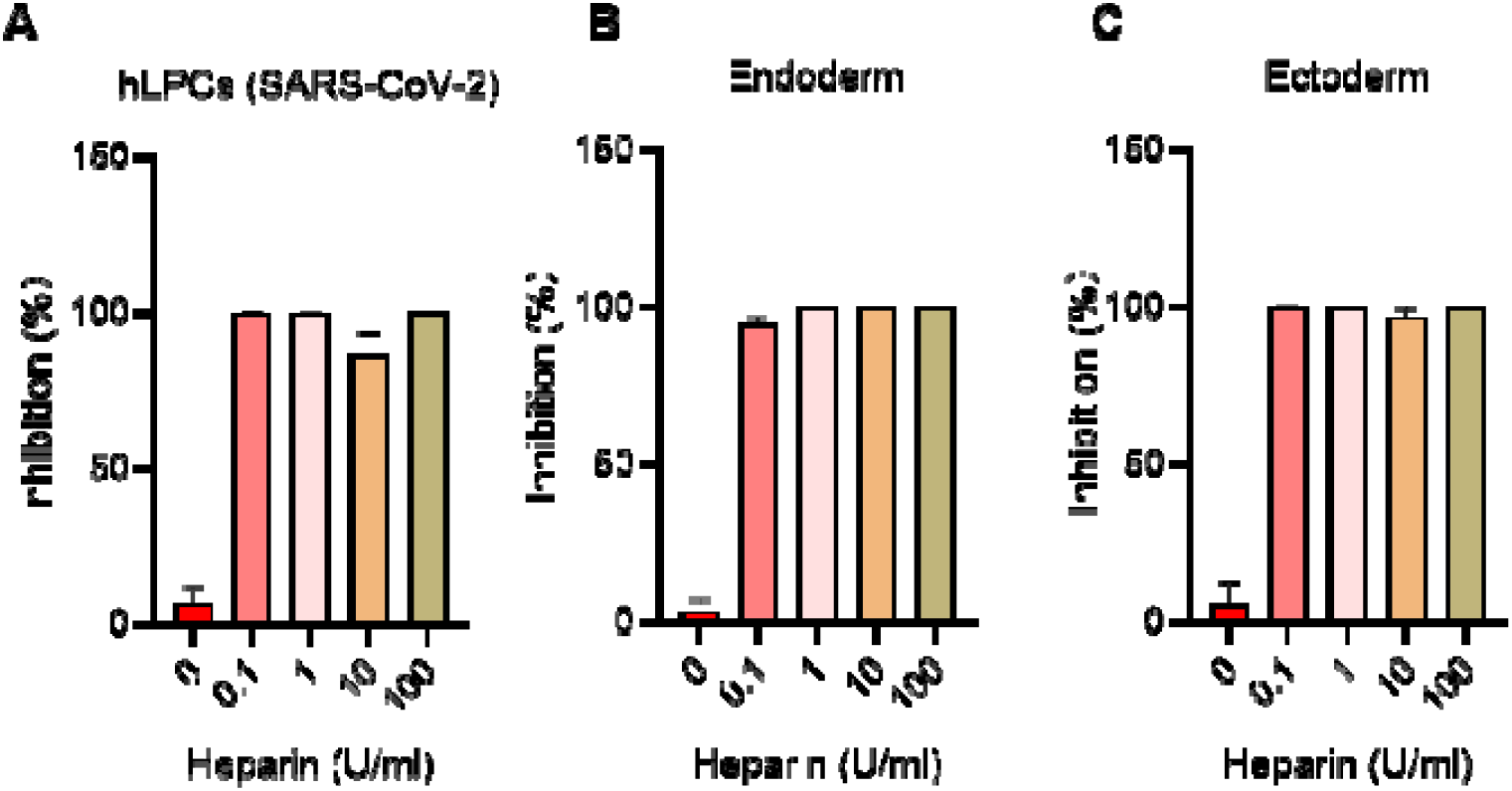
Inhibition of SARS-CoV-2 pseudovirus infection by heparin in hLPCs, endoderm and ectoderm cells. (**A**) hLPCs, (**B**) endoderm and (**C**) ectoderm were infected by SARS-CoV-2 pseudovirus at viral inocula of 650 TCID_50_/well with or without different concentration of heparin. hLPCs, human lung progenitor cells.

